# Conserved RNA-binding protein interactions mediate syntologous lncRNA functions

**DOI:** 10.1101/2024.08.21.605776

**Authors:** Xavier Sabaté-Cadenas, Perrine Lavalou, Caroline Jane Ross, Lee Chen, Dina Zielinski, Sophie Vacher, Mireille Ledevin, Thibaut Larcher, Matthieu Petitjean, Louise Damy, Nicolas Servant, Ivan Bièche, Igor Ulitsky, Alena Shkumatava

## Abstract

Syntologous long noncoding RNAs (lncRNAs) are loci with conserved genomic positions that often show little or no sequence similarity. Despite diverging primary sequences, lncRNA syntologs from distant species can carry out similar functions. However, determinants underlying conserved functions of syntologous lncRNA transcripts with no sequence similarity remain unknown. Using *CASC15* and melanoma formation as a paradigm for fast evolving lncRNAs and their functions, we found that human and zebrafish *CASC15* syntologs with no detectable sequence similarity retained their function across 450 million years of evolution. Similar to the *casc15*-deficient zebrafish, *CASC15-*mutant human melanoma cells show increased cell migration. Expression of human *CASC15* in zebrafish rescues loss of *casc15* function by attenuating melanoma formation. This conserved function is supported by a set of RNA-binding proteins, interacting with both zebrafish and human *CASC15* transcripts. Together, our findings demonstrate that conserved RNA-protein interactions can define functions of rapidly evolving lncRNA transcripts.

## INTRODUCTION

The identification of determinants underlying functions of long noncoding RNAs (lncRNAs) is key to understanding how cellular and molecular processes are modulated by these regulatory RNAs. Whereas several lncRNAs have been established as crucial players in various physiological and pathological processes, their functional domains and the determinants that define their molecular modes of action remain elusive^1–4^.

Analysis of evolutionary constraints on the primary sequence have been successfully used to identify functional elements within lncRNA transcripts^5–9^. However, unlike protein-coding genes, lncRNA sequences undergo fast evolutionary turnover and as such rarely exhibit longer stretches of sequences conserved throughout substantial evolutionary distances^8,10,11^. Whereas most protein-coding genes are well conserved, only ∼100 lncRNAs show linear sequence conservation between mammals and fish based on whole-genome alignments^8,10^. Alignment-based lncRNA sequence comparisons revealed that conserved sequences are usually restricted to relatively short stretches of 50-300 nucleotides that are embedded in rapidly evolving surrounding sequences^8,10^. This overall low evolutionary conservation of lncRNA sequences has hampered the discovery of orthologous genes in different vertebrate species which precludes defining functional elements within these transcripts.

In contrast to the low selective constraints on the primary lncRNA sequences, lncRNA genomic positions with respect to flanking orthologous protein-coding genes are often well preserved throughout vertebrate evolution^8,10^. Synteny has facilitated the identification of lncRNA transcriptional homologs (also referred as syntologs) with diverging primary sequences but conserved biological functions. One prominent example is the well-characterised *Xist* lncRNA that exhibits a high rate of sequence turnover between human and rodents^12^. Despite the rapidly evolving primary sequence, the syntologous human and mouse *Xist* transcripts are functionally conserved regulating gene dosage compensation in mammals^13,14^. Similarly, utilising synteny, 47 *roX* lncRNAs were identified in various Drosophilid species in addition to those previously described^15^. Despite their diverging sequences, transgenic expression of *roX* syntologs rescued male lethality in *roX*-null *D. melanogaster* caused by deficits in gene dosage compensation^15^. Thus, comparative analyses of syntologous lncRNAs present a powerful strategy for finding functionally related lncRNA transcripts that often evade detection based solely on sequence alignments.

Another approach to identify homologous lncRNAs and potentially define their functional sequence elements is through the comparison of short sequence motifs scattered throughout lncRNA transcripts^16^. Several algorithms have been developed to identify short conserved sequence elements and their combinations within RNA transcripts^17^. SEEKR groups lncRNAs with potentially related functions based on the similar content of the short sequence motifs called *k*-mers, which are defined as all possible combinations of nucleotides where *k* specifies the length of the motif^18–20^. Whereas SEEKR does not consider the relative order of *k*-mers within potentially functionally related lncRNAs with divergent sequences, LncLOOM identifies combinations of short motifs positioned in the same order within transcriptional homologs from different species^21^. Regardless of the method used for sequence analyses, the short RNA motifs usually correspond to binding sites of RNA-binding proteins (RBPs)^22^. As RNA-binding proteins are the main executors of lncRNA functions in the cell, identification of lncRNA-protein interactions is key to uncovering molecular mechanism of lncRNA action. Notably, only a fraction of RBPs binds to defined consensus motifs, whereas most RNA-binding proteins have limited sequence specificity^22,23^. The protein-RNA binding specificity is dictated not only by RNA recognition motifs but also by additional features such as local RNA secondary structure, flanking nucleotide composition, and/or bipartite motifs^22^.

The human lncRNA *CASC15* (*CAncer Susceptibility Candidate 15*) is a versatile molecule that has been reported to be a cancer-related lncRNA due to its association with multiple human carcinogenesis and tumorigenesis events^24–28^. Dependent on the cancer type and the cellular system, *CASC15* has been reported to act either as a tumour suppressor or oncogene^24–27^. Among reported cancer-linked functions, *CASC15*, which is frequently upregulated in melanoma tumours, has been shown to contribute to metastatic melanoma cell transition between proliferative and invasive states^25^. While *CASC15* has emerged as a cancer-associated lncRNA, the molecular mechanism of *CASC15* action, including its functional elements, remains poorly understood. Using *CASC15* as a paradigm lncRNA and melanoma formation as a functional read-out, here we show that syntologous lncRNA transcripts with no detectable sequence similarity attenuate melanoma formation by acting through a conserved repertoire of lncRNA-protein interactions. We demonstrate that lncRNAs can tolerate major changes in gene architecture, such as the length of the mature sequence and number of exons, while retaining their conserved biological functions that are defined by lncRNA-protein interactomes. Collectively, we show that conserved lncRNA-protein interactions can regulate critical cellular processes.

## RESULTS

### The lncRNA *CASC15* shows synteny but no sequence conservation across vertebrates

The lncRNA *CASC15* is located between the *SOX4 (SRY BOX 4*) and *PRL* (*PROLACTIN*) protein-coding genes (**Fig. 1a**). Examination of chromosome regions surrounding the *SOX4* loci in different vertebrates revealed the presence of synteny blocks with homologous genes that share a common gene order and transcriptional orientation (**Supplementary Fig. 1a**). Based on their location within the *SOX4* synteny block, we identified transcriptional homologs of *CASC15* in all analysed vertebrates including zebrafish (**Fig. 1a; Supplementary Fig. 1a**). Whereas *SOX4*–lncRNA synteny is highly preserved across vertebrates, BLASTN detected only low level of sequence similarity at the splice junction of exon 2 in tetrapods and mammals (**Supplementary Fig. 1b**). Beyond synteny, no sign of *CASC15* sequence or gene structure similarities could be found in more distant vertebrates such as zebrafish. The only exception was a short, highly conserved region at the 5’ end of the *CASC15* gene (**Fig. 1a; Supplementary Fig. 1c).** This sequence homology likely results from the conservation of *cis*-regulatory elements in the promoter region of *CASC15* as reported for other syntologous lncRNAs that often show sequence conservation only in promoter-proximal regions^10,11^. In addition to synteny, *CASC15* lncRNA transcripts from distant species, such as humans and zebrafish, share similar expression patterns across adult tissues and organs, with the highest expression detected in the ovary^29^ (**Fig. 1b, c**). The conserved tissue expression pattern and synteny suggest that *CASC15* transcripts have retained regulatory programs and potentially conserved functions across vertebrates.

**Figure 1:**
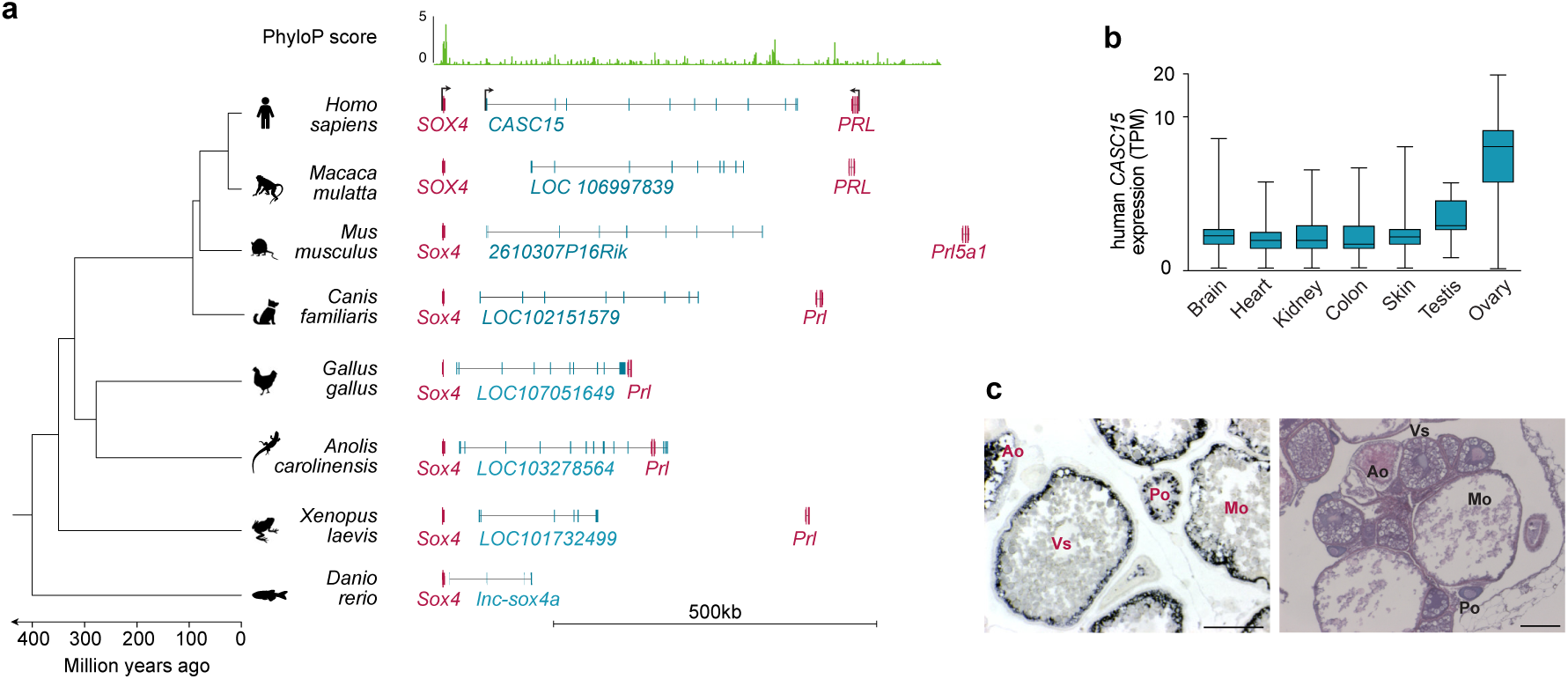
Synteny and conserved expression pattern of the lncRNA *CASC15*. **(a)** Genomic locus of *CASC15* in different vertebrates. Representative isoforms of *CASC15* are shown. Blue boxes represent *CASC15* exons, red boxes represent *SOX4*-coding exons and red boxes represent *PRL*-coding exons. Vertebrate sequence conservation is measured PhyloP scores derived from 100-way vertebrate multiple alignment and shown in green. **(b)** Expression of human *CASC15* across different organs from the Genotype-Tissue Expression (GTEx) project. TPM, transcripts per million. **(c)** *casc15* expression by *in situ* hybridization on adult zebrafish ovary sections (left panel). Haematoxylin and eosin staining of zebrafish ovary sections (right panel). Primary oocytes (Po); Vitellogenic stage (Vs); Mature oocyte (Mo); Atretic oocyte (Ao). Scale bars represent 200 µm.

### The zebrafish lncRNA *casc15* inhibits melanoma formation *in vivo*

Human *CASC15* is elevated in multiple human cancers including primary and metastatic melanomas (**Supplementary Fig. 2a**; **Fig. 2a**). Furthermore, *CASC15* regulates melanoma progression by controlling the switch between proliferative and invasive states of human melanoma cells^25^. Thus, we sought to examine whether the regulatory function of *CASC15* in melanoma formation is retained in distant syntologs. To test whether, similar to human *CASC15*, zebrafish *casc15* regulates melanoma formation *in vivo*, we used a zebrafish melanoma system that employs expression of the human oncogenic NRAS^G12D^ protein driven by the melanocyte-specific *mitfa* promoter^30,31^. We injected zebrafish embryos at the one-cell stage with a construct expressing *mitfa*:*NRAS^G12D^*and monitored melanoma formation for several weeks post-injection (**Supplementary Fig. 2b, c**). Animals expressing NRAS^G12D^ were evaluated for visible pigmentation defects and tumour formation^30,31^ (**Supplementary Fig. 2d**). First, we tested *casc15* expression in the NRAS^G12D^-induced zebrafish tumours. Similar to *CASC15* upregulation in human melanoma samples, the zebrafish *casc15* transcript was significantly elevated in tumours of NRAS^G12D^-injected animals as compared to the skin of un-injected fish suggesting activation of similar gene expression programs during zebrafish and human melanoma formation (**Figure 2a, b)**. To test the function of zebrafish *casc15*, we used our previously generated *lnc-sox4a^ΔTSS^*zebrafish allele, hereafter referred to as *casc15^ΔTSS^*, which annulled *casc15* expression in embryos and in the majority of adult tissues including skin^29^ (**Supplementary Fig. 2e**). When analysing different genotypes of zebrafish expressing NRAS^G12D^ driven by the *mitfa* promoter, a much greater and significantly higher percentage of *casc15^ΔTSS^*fish showed tumour formation as compared to wild-type or unrelated lncRNA *malat1^-/-^*mutant fish injected with the same *mitfa*:*NRAS^G12D^* construct^29^ (**Fig. 2c**). Moreover, pigmentation defects appeared and tumours reached more advanced tumour stages faster in *casc15^ΔTSS^*animals expressing the *mitfa*:*NRAS^G12D^* transgene than in wild-type fish expressing the same transgene (**Fig. 2d**). In addition, *casc15^ΔTSS^* fish expressing *mitfa*:*NRAS^G12D^* developed highly proliferative internal tumours (**Supplementary Fig. 2f-h**). Whereas the internal tumorigenesis was a rare event affecting only 1.3% of *casc15^ΔTSS^* fish expressing NRAS^G12D^ (2 out of 169 examined animals), no internal tumours were detected in any other examined genetic backgrounds expressing NRAS^G12D^ (0 out of 667 examined animals). In summary, our results demonstrate that the zebrafish *casc15* lncRNA attenuates melanoma onset and progression *in vivo*.

**Figure 2:**
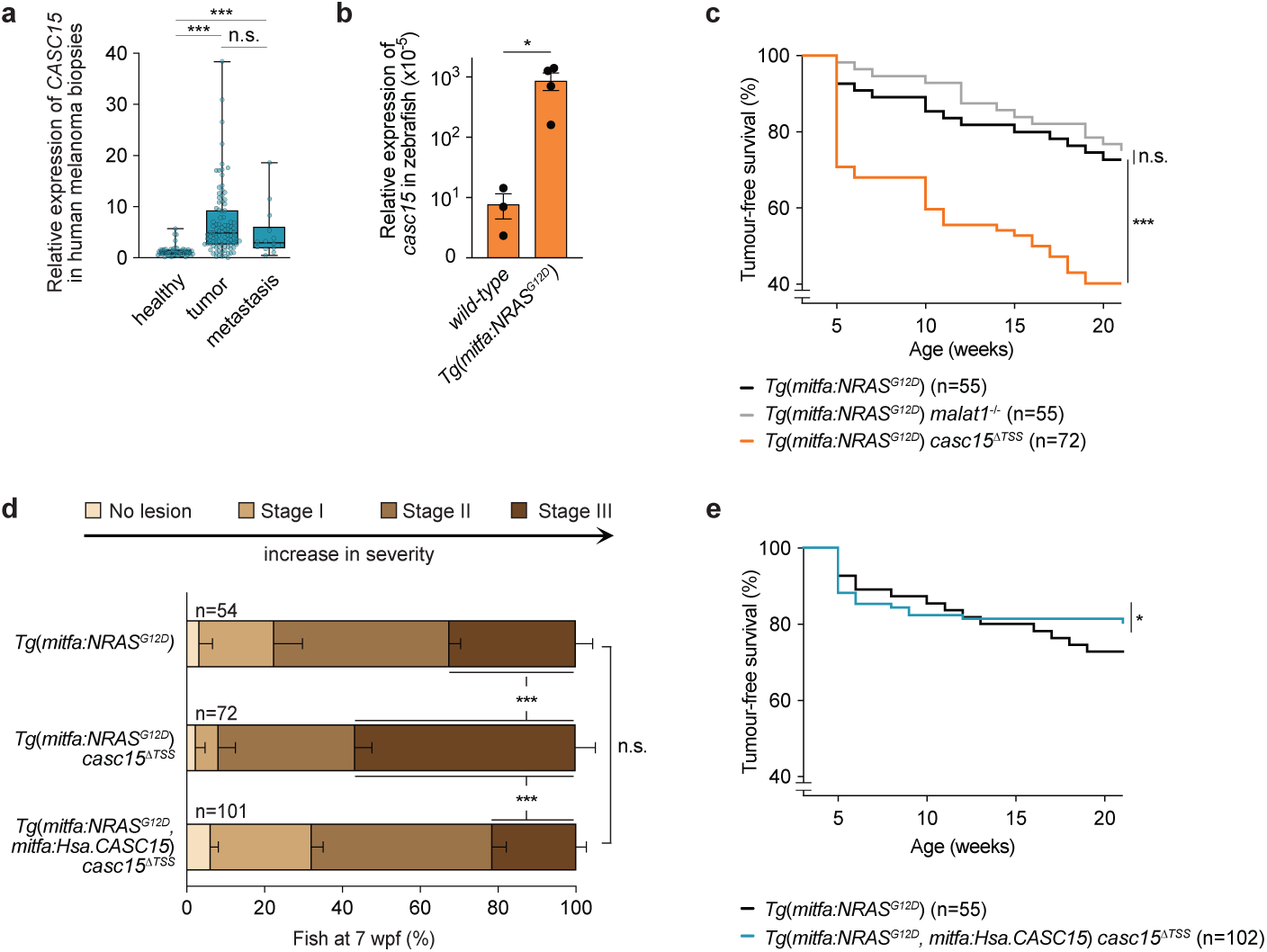
*casc15* attenuates melanoma formation and progression in zebrafish. **(a)** qRT-PCR analysis of human *CASC15* expression in patient melanoma biopsies (healthy, tumour and metastatic tissues). *TBP* was used as a reference gene; Mann-Whitney test: \*\*\**P* < 0.001; not significant (ns). **(b)** qRT-PCR analysis of *casc15* expression in the wild-type zebrafish skin and melanoma tumours isolated from zebrafish expressing the NRAS^G12D^ driven by the *mitfa* promoter. *eef1α1* was used as a reference gene. Data are presented as mean ± s.e.m.; unpaired t-test: \**P* < 0.05 **(c)** Kaplan–Meier plot of melanoma-free period in wild-type animals compared to *malat1*^-/-^ and *casc15^ΔTSS^* mutant zebrafish, injected with NRAS^G12D^. Data are analysed using Mantel-Cox test: \*\*\**P* < 0.001; not significant (ns). **(d)** Percentage of wild-type and *casc15^ΔTSS^* zebrafish expressing NRAS^G12D^ driven by the *mitfa* promoter exhibiting different melanoma stages at seven weeks post-fertilization (wpf). Data are presented as mean ± s.e.m.; two-way ANOVA with Tukey’s multiple comparisons test; \*\*\**P* < 0.001, not significant (ns). **(e)** Kaplan–Meier plot of melanoma-free period in wild-type zebrafish expressing NRAS^G12D^ driven by the *mitfa* promoter and *casc15^ΔTSS^*zebrafish co-expressing NRAS^G12D^ and human *CASC15* (*Hsa.CASC15*) driven by the *mitfa* promoter. Data are analysed using Mantel-Cox tests: \**P* < 0.05.

### Expression of human *CASC15* in zebrafish melanocytes slows down melanoma progression

We carried out inter-species genetic complementation experiments to examine whether zebrafish and human *CASC15* transcripts have equivalent functions during melanoma formation *in vivo*. We co-expressed the mature human *CASC15* transcript (*Hsa.CASC15*) together with NRAS^G12D^ in melanocytes of *casc15^ΔTSS^* zebrafish and tested melanoma formation in these animals (**Supplementary Fig. 2i**). Melanocyte-specific expression of human *CASC15* together with NRAS^G12D^ in *casc15^ΔTSS^* zebrafish significantly decreased the percentage of animals developing tumours and was comparable to wild-type fish expressing NRAS^G12D^ (**Fig. 2e**). In addition, *Tg(mitfa:*NRAS^G12D^, *mitfa:Hsa.CASC15*) *casc15^ΔTSS^* animals exhibited reduced melanoma formation and tumour progression, which were comparable to wild-type animals expressing NRAS^G12D^ (**Fig. 2d**). By contrast, co-expression of NRAS^G12D^ together with unrelated *Venus*, which served as a rescue specificity control, in melanocytes of *casc15^ΔTSS^* fish did not affect melanoma formation (**Supplementary Fig. 2 j**). Notably, the rescue effect of the human *CASC15* transcript in *Tg(mitfa:*NRAS^G12D^, *mitfa:Hsa.CASC15*) *casc15^ΔTSS^* fish was not due to the decreased expression levels of the *mitfa:*NRAS^G12D^ transgene (**Supplementary Fig. 2k**). Together, these results suggest that despite diverging sequences, *CASC15* syntologs from distant vertebrate species carry out conserved functions to repress melanoma formation *in vivo*.

### *CASC15* attenuates human melanoma cell migration

As siRNA-based depletion of *CASC15* has been previously reported to regulate human melanoma cell phenotype switching between proliferative and invasive states^25^, we sought to further test the regulatory function of *CASC15* in human melanoma cells by generating *CASC15* mutant alleles. We profiled several melanoma cell lines for *CASC15* expression and chose 501mel cells based on the high levels of *CASC15* (**Supplementary Fig. 3a)**. In 501mel cells, the main *CASC15* isoform is produced from a downstream transcriptional start site (TSS2) generating a 1.5 kb transcript comprised of four exons (exon 9b - exon 12) (**Fig. 3a**). Expression of similar short *CASC15* isoforms has been previously reported in other cancer cell lines including YDFR.SB3, WP and RKTJ-CB1 melanoma, and SH-SY5Y and SK-N-SH neuroblastoma cells^24–26^. To generate loss of *CASC15* function, we engineered homozygous *CASC15^Δexon^*^12^ 501mel cells where the largest terminal exon (exon 12) which encodes 1.15 kb of the *CASC15* transcript is deleted (**Fig. 3a, Supplementary Fig. 3b, c**). Upon removal of exon 12, a residual *CASC15* transcript was detectable in *CASC15^Δexon^*^12^ 501mel cells albeit at ∼40% of wild type *CASC15* levels (**Fig. 3a, Supplementary Fig. 3d**). In summary, the *CASC15^Δexon^*^12^ allele encodes a transcript missing ∼1150 nucleotides that is expressed at markedly lower levels.

**Figure 3:**
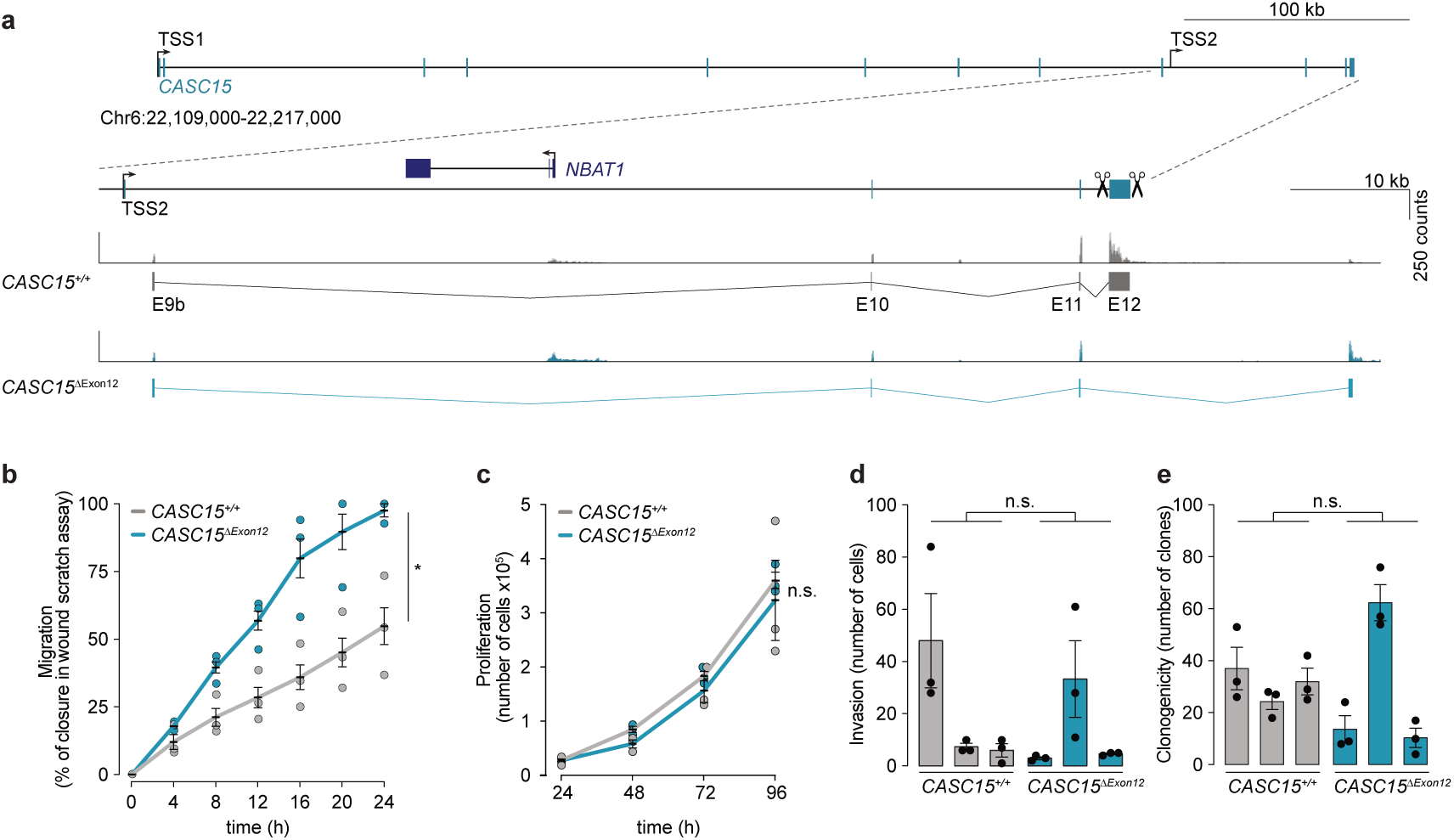
*CASC15* alleviates cell migration in human melanoma cells. **(a)** Generation of the *CASC15^Δexon^*^12^ allele in 501mel cells. Top panel, the *CASC15* locus. Dotted line delineates a zoom-in of the 3′ region of *CASC15*. Scissors indicate the location of gRNAs to target the largest 3′ exon for deletion. Bottom panel, RNA-Seq confirmed generation of the *CASC15^Δexon^*^12^ allele. RNA-Seq tracks are set to the same scale. Schematics of the *CASC15* transcript in unedited *CASC15^+/+^* and *CASC15^Δexon^*^12^ 501mel cells are presented below each RNA-Seq track. TSS, transcriptional start site. **(b)** Wound-healing assay quantification in 501mel cells. Three biological replicates (independently generated cell lines) are shown for each genotype. Data are presented as mean ± s.e.m.; two-way ANOVA test: \**P* < 0.05. **(c)** Proliferation assay quantification. Three biological replicates (independently generated cell lines) are shown for each genotype. Data are presented as mean ± s.e.m.; two-way ANOVA test: not significant (ns). **(d)** Invasion assay quantification. Three biological replicates (independently generated cell lines) are shown for each genotype. Data are presented as mean ± s.e.m.; unpaired t-tests: not significant (ns). **(e)** Clonogenicity assay quantification. Three biological replicates (independently generated cell lines) are shown for each genotype. Data are presented as mean ± s.e.m.; unpaired t-tests: not significant (ns).

Next, we subjected *CASC15^Δexon^*^12^ 501mel cells to a series of cellular assays. In a wound-healing assay, *CASC15^Δexon^*^12^ 501mel cells showed a significantly increased cell migration when compared to *CASC15^+/+^* 501mel cells (**Fig. 3b**). By contrast, no significant changes in cell proliferation, invasiveness or clonogenicity were detected (**Fig. 3c-e; Supplementary Fig. 3e-g**). Consistent with the increased migration of *CASC15^Δexon^*^12^ cells, Gene Ontology (GO) analyses of transcriptome changes showed that genes upregulated in *CASC15^Δexon^*^12^ 501mel cells were enriched for functions associated with angiogenesis, cell migration and taxis (**Supplementary Fig. 3h-j**). By contrast, genes downregulated in *CASC15^Δexon^*^12^ 501mel cells were not enriched for any GO terms (**Supplementary Fig. 3i**). Taken together, our results demonstrate that similar to the *in vivo* zebrafish melanoma model, *CASC15* attenuates cell migration of human melanoma cells.

### Syntologous *CASC15* transcripts differ in their predicted RNA motif contents

We sought to determine the molecular rationale underlying the conserved functions of human and zebrafish *CASC15* transcripts that lack detectable sequence homology. As many lncRNAs modulate gene expression *in cis*^32^ and *CASC15* has been reported to regulate its neighbouring protein-coding gene *SOX4* in RUNX1-rearranged acute leukaemia^27^, we first quantified the levels of *Sox4a* (the zebrafish *SOX4* homolog that is genomically adjacent to *casc15*) in zebrafish NRAS^G12D^-induced tumours and *SOX4* in human 501mel cells, respectively. No significant changes in the *Sox4a* mRNA levels were detected in tumours isolated from *Tg(mitfa:*NRAS^G12D^) *casc15^ΔTSS^* fish as compared to tumours isolated from wild-type animals expressing the *mitfa:*N*RAS^G12D^*transgene (**Fig. 4a**). Similarly, *SOX4* mRNA expression was unchanged in *CASC15^Δexon^*^12^ 501mel cells as compared to *SOX4* levels in unedited 501mel cells (**Fig. 4b**) indicating that in the specific melanoma context, the function of *CASC15* is independent of regulation of *SOX4*.

**Figure 4:**
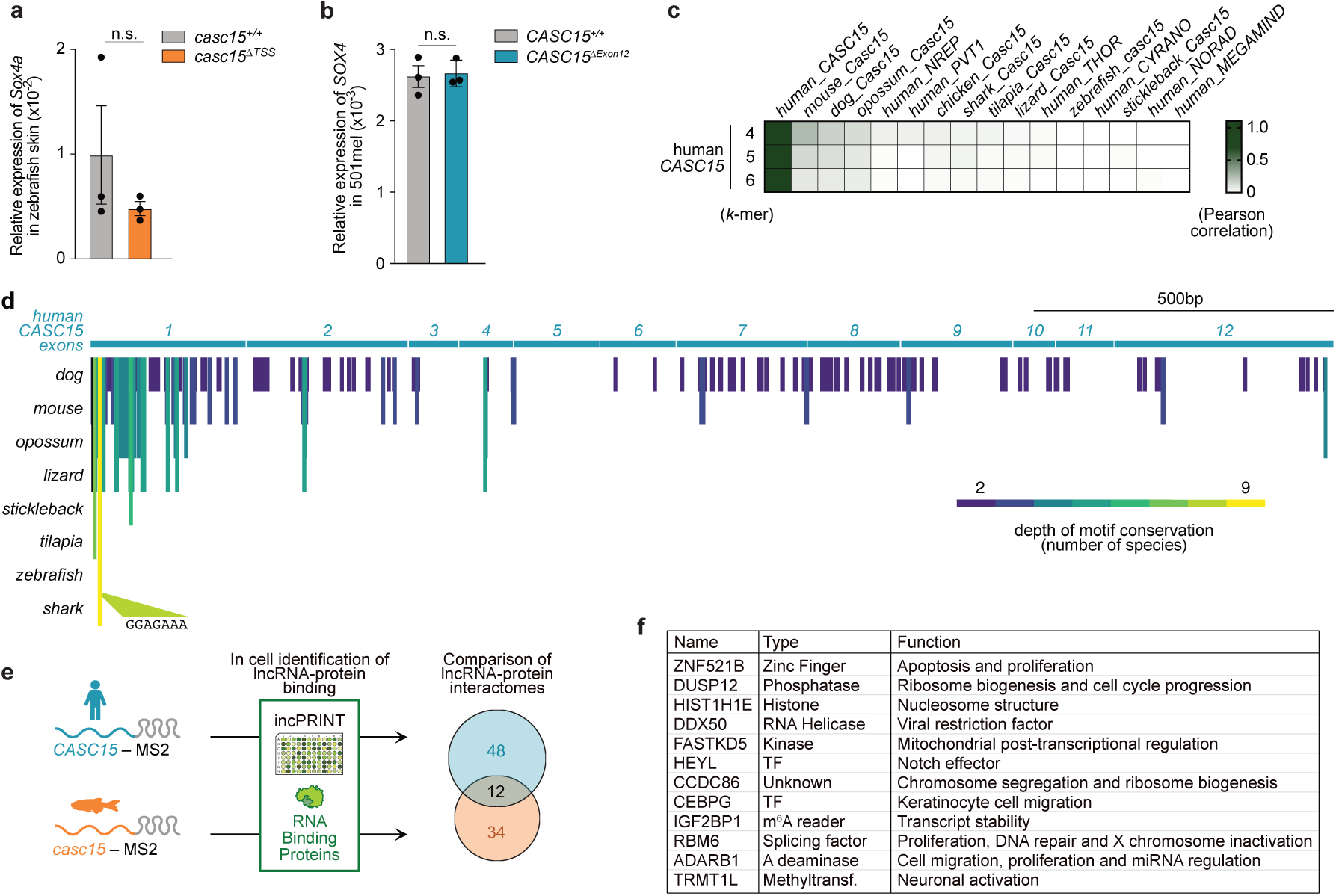
Human and zebrafish *CASC15* transcripts exhibit different composition of RNA motifs but share a set of protein interactors. **(a)** qRT-PCR analysis of *sox4a* expression in melanoma tumours isolated from wild-type and *casc15^ΔTSS^*zebrafish expressing NRAS^G12D^ driven by the *mitfa* promoter. Three biological replicates are shown for each genotype. *eef1α1* was used as a reference gene. Data are presented as mean ± s.e.m.; unpaired t-test: not significant (ns). **(b)** qRT-PCR measuring *SOX4* mRNA expression levels in 501mel cells. Three biological replicates (independently generated cell lines) are shown for each genotype. *ß-ACTIN* was used as a reference gene. Data are presented as mean ± s.e.m.; unpaired t-test: not significant (ns). **(c)** *k*-mer content analysis between human *CASC15* and the indicated vertebrate orthologs and unrelated human lncRNAs using SEEKR algorithm. *k*-mer length is indicated in the *y*-axis. All human lncRNAs were used as normalization set. **(d)** Distribution of short RNA motifs in exons of *CASC15* identified by the LncLOOM algorithm to be conserved among vertebrates. Identified elements are represented by vertical bars color-coded by conservation depth. Vertebrate species containing the motif are indicated in the left. The human *CASC15* transcript was used as the anchor. **(e)** Experimental strategy to identify proteins interacting either with zebrafish *casc15* or human *CASC15*. The identification of lncRNA interactomes was performed by incPRINT using MS2-tagged human and zebrafish bait RNAs. The library of ∼3000 FLAG-tagged human proteins was interrogated for binding either with human or zebrafish *CASC15* transcripts. Venn diagram represents comparison of human and zebrafish *CASC15* interactomes and indicates number of shared proteins.

Next, we tested if the conserved function of the syntologous *CASC15* transcripts can be supported by a set of short, conserved RNA motifs (*k*-mers) that remained undetected by BLASTN as shown for some other lncRNAs^18,21^. We used two different algorithms to define conserved motif elements within *CASC15* transcriptional homologs. First, we compared the *k*-mer content of the human *CASC15* to other vertebrate *CASC15* transcriptional homologs using the SEEKR algorithm^18^. We found that the *k*-mer content of *CASC15* differs substantially among examined vertebrates (**Fig. 4c; Supplementary Fig. 4a**). For instance, the 4-mer profile of human *CASC15* was more similar to other unrelated human lncRNAs (a Pearson’s R=0.07 for *NREP* and *PVT1*) than to zebrafish *casc15* (a Pearson’s R=0.01). (**Fig. 4c)**. Second, we used the graph-based algorithm LncLOOM that identifies conserved ordered RNA motifs within syntenic lncRNAs^21^. Using LncLOOM, only one 7-mer conserved between zebrafish and human was identified at the 5′ end of the *CASC15* transcripts (**Fig. 4d**). To avoid potential interference of the deeply conserved sequences in the first exon of *CASC15* (**Supplementary Fig. 1b**) with the search of conserved sequence motifs distributed along the transcript, we excluded the first exon from the LncLOOM analysis. No motifs were found to be conserved to zebrafish (**Supplementary Fig. 4b)**. Altogether, currently available algorithms identified no conserved RNA motifs within human and zebrafish *CASC15* transcriptional homologs. Our analyses suggest that the functional conservation of human and zebrafish *CASC15* transcripts is not supported by detectable sequence homology nor can it be explained by the regulation of its protein-coding neighbour gene *SOX4*.

### Human and zebrafish *CASC15* transcripts share a set of RNA-binding protein interactors

As vertebrate *CASC15* transcripts show neither longer stretches of RNA sequence homology nor detectable microhomology, we sought to examine whether other determinants define the conserved function of *CASC15* transcripts. As RNA-binding proteins commonly bind cryptic motifs with low compositional complexity^22^, we reasoned that functionally related lncRNAs with divergent sequences may interact with the conserved sets of RNA-binding proteins defining similar cellular and molecular functions of lncRNAs. To determine proteins interacting with human and zebrafish *CASC15* transcripts, we employed incPRINT, a technology to comprehensively identify RNA-protein interactions^33^. We used full-length, mature human *CASC15* (∼2 kb) and zebrafish *casc15* (∼1,8 kb) RNAs tagged with MS2 repeats as bait RNAs to interrogate our library of ∼3000 FLAG-tagged human proteins that includes most known RNA-binding proteins, chromatin-associated proteins and transcription factors^33^ (**Fig. 4e**). Notably, zebrafish and human *CASC15* RNAs used for incPRINT did not include the 5′ conserved sequence (**Supplementary Fig. 1c)**. We identified 48 and 34 proteins that interacted with either human or zebrafish *CASC15* RNAs, respectively (**Supplementary Table 1**). Twelve of the identified proteins were shared between human and zebrafish *CASC15* interactomes (**Fig. 4e, f**). Thus, our results suggest that *CASC15* transcripts that lack detectable sequence homology bind a shared set of specific protein interactors providing a rationale for the conserved regulatory function of *CASC15*.

### DUSPl2 and DDX50 mediate *CASC15* function

To identify proteins that are key for the *CASC15* function, we individually depleted each of the 12 shared *CASC15*-interacting proteins in 501mel cells using esiRNAs^34^ and subsequently tested cell migration in the wound-healing assay (**Fig. 5a**). Depletion of DDX50 and DUSP12 specifically increased cell migration but had no effect on cell proliferation, thus matching cellular features of the *CASC15^Δexon^*^12^ 501mel cell lines (**Fig. 5b-d; Supplementary Fig. 5a, b**). By contrast, the depletion of the other ten identified *CASC15*-interacting proteins did not affect the migration of 501mel cells in the wound-healing assay (**Fig. 5b)**.

**Figure 5:**
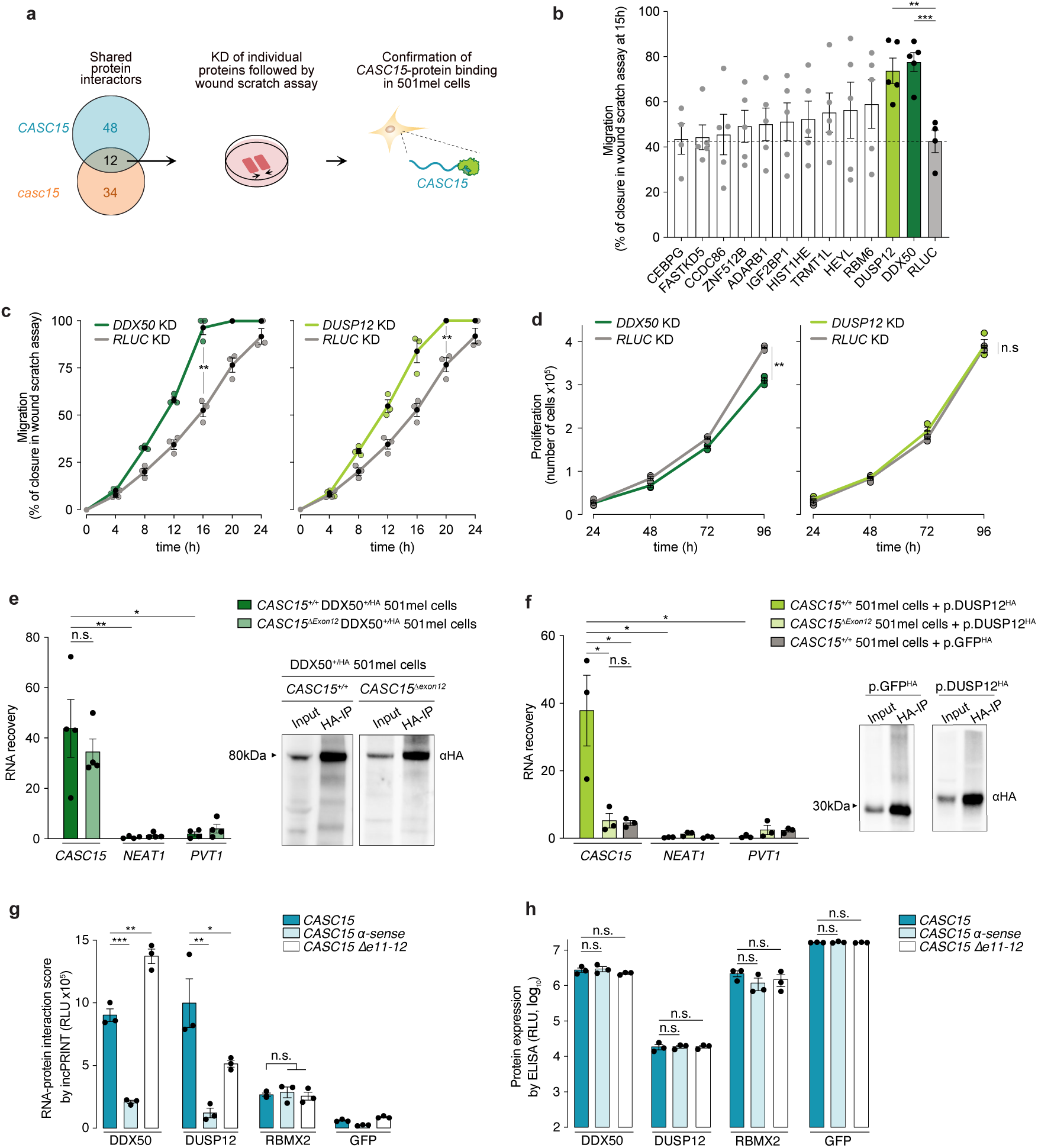
DDX50 and DUSP12 are key executers of the *CASC15* function. **(a)** Experimental strategy to identify functionally important proteins. **(b)** Wound-healing assay in 501mel cells upon depletion of the indicated proteins. Data represent percentage of closure at 15h. *RLUC* was used as negative control. Data from at least four independent experiments is presented as mean ± s.e.m.; unpaired t-test: \*\**P* < 0.01; *** *P* < 0.001. **(c)** Wound-healing assay in 501mel cells upon depletion of either DDX50 (left panel) or DUSP12 (right panel) over the 24h-period. *RLUC* was used as negative control. Data from three independent transfections is presented as mean ± s.e.m.; two-way ANOVA test: \*\**P* < 0.01. **(d)** as in (c), for the proliferation assay. Data from three independent transfections is presented as mean ± s.e.m.; two-way ANOVA test: \*\**P* < 0.01; not significant (ns). **(e)** RNA immunoprecipitation (RIP) of the endogenously HA-tagged DDX50 protein either in *CASC15^+/+^* or *CASC15^Δexon^*^12^ 501mel cells. Left panel, RNA recoveries indicate RNA expression levels of the indicated transcripts in the immunoprecipitated eluates, normalised to *ß-ACTIN* mRNA and to input samples. Each RIP experiment was performed in four independent biological replicates. Data are presented as mean ± s.e.m.; unpaired t-tests: \**P* < 0.05; \*\**P* < 0.01; not significant (ns). Right panel, Western blot for HA-DDX50 using anti-HA antibody on samples isolated from either *CASC15^+/+^*or *CASC15^Δexon^*^12^ 501mel cells. **(f)** RNA immunoprecipitation (RIP) of the HA-tagged DUSP12 and HA-tagged GFP proteins expressed from plasmids either in *CASC15^+/+^* or *CASC15^Δexon^*^12^ 501mel cells. Left panel, as in **(e)**. Right panel, Western blot for HA-GFP and HA-DDX50 using anti-HA antibody on samples isolated from *CASC15^+/+^*501mel cells. **(g)** Interaction intensities between indicated *MS2*-tagged RNAs and DDX50, DUSP12, RBMX2 and GFP measured by incPRINT. The non-discriminatory RBMX2 protein was used to control for RNA expression levels. GFP was used as a negative control. Data from three biological replicates are presented as mean ± s.e.m.; unpaired t-tests: \**P* < 0.05; \*\**P* < 0.01; \*\*\**P* < 0.001; not significant (ns). RLU, Relative Luminescence Units. **(h)** Protein expression levels measured by ELISA in the incPRINT assay. Data from three biological replicates are presented as mean ± s.e.m.; unpaired t-tests: \*\**P* < 0.01; not significant (ns). RLU, Relative Luminescence Units.

To confirm the interaction of DDX50 and DUSP12 with the *CASC15* lncRNA in 501mel cells and to identify *CASC15* sequence elements required for DDX50 or DUSP12 binding, we performed a series of experiments in which we conducted RNA immunoprecipitation (RIP) after UV crosslinking. First, we engineered *DDX50^+/HA^* and *DUSP12^+/HA^* alleles in 501mel cells. The sequence encoding the HA epitope tag has been inserted into either the *DDX50* or *DUSP12* loci generating an endogenously expressed *N*-terminally epitope-tagged HA-DDX50 or HA-DUSP12, respectively (**Supplementary Fig. 5c, d**). RIP experiments after UV-cross-linking of cells followed by qRT-PCR analyses identified a significant and specific association of the *CASC15* transcript with DDX50, confirming their interaction in 501mel cells (**Fig. 5e**). The same significant enrichment of *CASC15* was detected when the UV-RIP experiment was performed in *CASC15^Δexon^*^12^, *DDX50^+/HA^* 501mel cells suggesting that DDX50 also binds to the truncated *CASC15^Δexon^*^12^ transcript (**Fig. 5e**). In contrast to endogenously tagged HA-DDX50, we were not able to detect HA-tagged DUSP12 protein in *DUSP12^+/HA^* 501mel cells (**Supplementary Fig. 5e**). Using a commercially available antibody, we specifically detected endogenous DUSP12 protein in 501mel cells (**Supplementary Fig. 5b**). However, using this antibody, we were unable to enrich the DUSP12 protein after the IP. Thus, we expressed HA-DUSP12 from a plasmid transfected into *CASC15^+/+^* and *CASC15^Δexon^*^12^ 501mel cells and proceeded with the UV-RIP experiment followed by qRT-PCR. A significant and specific enrichment of the endogenous *CASC15* transcript was detected with HA-DUSP12 confirming their interaction in unedited *CASC15^+/+^* 501mel cells (**Fig. 5b**). By contrast, no enrichment of the endogenous *CASC15^Δexon^*^12^ transcript was detected with the HA-DUSP12 protein (**Fig. 5f**) indicating that the sequence of exon 12 is required for the DUSP12-*CASC15* interaction. Consistent with the RIP results, incPRINT experiments confirmed that the interaction between truncated *CASC15* RNA and DDX50 protein remained unaffected and was comparable to full-length *CASC15* (**Fig. 5g; Supplementary Fig. 5d**). By contrast, *CASC15* RNA-DUSP12 interaction was diminished in the absence of exon 12 as compared to full-length *CASC15* corroborating the results of the RIP experiments (**Fig. 5g**). Notably, DUSP12 and DDX50 expression levels remained unchanged with all tested *CASC15* bait RNAs (**Fig. 5h**). Together, our data suggest that the regulatory function of *CASC15* is mediated by its interaction with two key proteins, the DDX50 RNA helicase and the DUSP12 phosphatase. Importantly, we identified exon 12 as a functional sequence element of human *CASC15* required for the specific interaction with DUSP12.

## DISCUSSION

Using syntenic conservation, we found transcriptional homologs of the lncRNA *CASC15* in all examined vertebrates, including some of the most distal vertebrates in the vertebrate phylogeny. Comparative analyses of syntologous *CASC15* transcripts from distant vertebrate species confirmed the rapid rate of lncRNA sequence turnover. Whereas a very low level of sequence conservation was found at a few exon-intron junctions of *CASC15* in reptiles and tetrapods, there was no detectable sequence similarity between mammals and species as evolutionarily distant as zebrafish. Nonetheless, our genetic complementation experiments demonstrated that human and zebrafish syntologs carry out conserved *CASC15* functions. Transgenic expression of human *CASC15* in the zebrafish *casc15* mutant rescued accelerated melanoma formation suggesting that the regulatory role of *CASC15* is retained in the syntologs despite divergence in their sequence. The phenomenon of lncRNA functional conservation with very poor or no detectable primary sequence homology is not restricted to the case of *CASC15* and has been reported for other orthologous lncRNAs using complementation experiments in various biological and cellular systems^15,20,35,36^. Thus, our functional analyses of the *CASC15* syntologs provide further evidence that lncRNAs can tolerate extensive sequence changes without functional disruption.

What are the determinants of the conserved biological lncRNA functions that are not primarily defined by conserved sequence elements? For *CASC15* syntologs, we explored several possibilities, including RNA motif content, *cis*-acting regulation of a neighbouring protein-coding gene and conserved lncRNA-interacting proteomes. We found no evidence for the *cis*-acting regulation of the *SOX4* gene by *CASC15*; neither did we find any conserved sequence motifs within the *CASC15* transcripts or when compared to other lncRNAs. Remarkably, we found that the most distantly related human and zebrafish *CASC15* syntologs share a substantial set of interacting proteins. From the specific set of shared *CASC15* interactors, two proteins-DDX50 (an ATP-dependent RNA helicase of the DEAD-box family) and DUSP12 (a phosphatase with dual specificity for serine/threonine and tyrosine phosphoresidues) - are key for the *CASC15* function in the context of the cell migration in human melanoma cells. Whereas the precise role of DDX50 and DUSP12 in melanoma formation and how *CASC15* modulates it remain to be addressed, these two proteins hold promise for novel therapeutic strategies as they have been associated with human cancers^37–41^. Notably, the residual set of *CASC15*-protein interactions shared between human and zebrafish transcripts may be functionally important in other biological and/or cellular contexts. Alternatively, they may regulate melanoma cell migration in a redundant manner which cannot be addressed by the depletion of individual proteins^42^. Together, identification and comparison of protein interactomes of distantly related lncRNAs and their further functional interrogation presents a powerful strategy for understanding the principles of lncRNA functionality and evolution.

The concept of rapidly evolving lncRNAs acting as flexible protein binding platforms^43–45^ aligns with the idea that many RNA-binding proteins have limited sequence specificity and recognise low complexity sequence motifs, which are often composed of just one or two base types^22^. Such cryptic binding motifs are not easy to identify as they evade detection with currently available comparative algorithms that usually rely on primary sequence conservation with perfect matches of 10+ consecutive bases and are blind to relatively degenerate sequences recognized by most RNA binding proteins. As such, more functional RNA elements may be embedded within distantly related lncRNA syntologs with diverging sequences. Here, we identified the last exon of *CASC15* as its functional sequence, as its removal abrogates *CASC15* function, resulting in faster melanoma cell migration. While no conserved sequence motifs were detected within this *CASC15* region, we showed that DUSP12 binds specifically to this region of *CASC15*. By contrast, the binding sites of DDX50 are not exclusive to the last exon of *CASC15* but are likely scattered throughout other parts of the transcript, highlighting the diversity of protein interaction modalities within the same transcript. In summary, we demonstrated that by interacting with a conserved set of RNA-binding proteins, lncRNAs can retain similar cellular functions even in the absence of identifiable conserved sequence motifs.

## METHODS

### Zebrafish melanoma model

All zebrafish were bred and maintained at the Institut Curie, Paris using standard procedures^46^. Animal care and use for this study were performed in accordance with the recommendations of the European Community (2010/63/UE) for the care and use of laboratory animals. Experimental procedures were specifically approved by the ethics committee of the Institut Curie CEEA-IC (Authorization APAFIS#13883-201803021206135-v1 and APAFIS#28477-2020120112502053 v2 given by National Authority) in compliance with the international guidelines.

To express oncogenic human *NRAS^G12D^* in zebrafish melanocytes, pDEST-Tol2-*mitfa:NRAS^G12D^* plasmid (gift of Adam Hurlstone, University of Manchester) was used. For genetic complementation experiments, the pDEST-Tol2-*mitfa:NRAS^G12D^* plasmid was modified by inserting either *mitfa:Venus* or *mitfa:CASC15* into the ClaI site. Human *CASC15* was amplified from FirstChoice Human brain Total RNA (Ambion, #6050) using Gibson assembly (New England Biolab, #E2621). An artificial intronic sequence was inserted between exon 2 and exon 3 of the mature *CASC15* transcript to insure correct processing of the transcript. To generate Tol2 transposase mRNA, pCS-TP plasmid (Adam Hurlstone, University of Manchester) was digested by Not I and RNA synthesized using HiScribe SP6 RNA Synthesis Kit (NEB, #E2070) and RNA was purified with the RNeasy Mini Kit (QIAGEN, #74104). Zebrafish embryos were co-injected into the one-cell stage with 50 ng/µl Tol2 mRNA, 50 ng/µl of the respective plasmid and 0.01% PhenolRed. At least two independent rounds of injections were performed for each experiment.

Injected embryos were selected for *Venus* expression in the eye lens at 72 hpf. Starting from 5 to 21 weeks post-fertilization, melanoma-induced zebrafish were individually evaluated each week and classified according to their melanoma progression stage: No Lesion; Stage I = pigmentation defects or localized nevi; Stage II = extended nevi, Radial Growth Progression; Stage III = Tumour/Hyperplasia; Vertical Growth Progression^47^ (**Supplementary Fig. 2e**). No further randomization or blinding was applied during data acquisition and analysis. All animals presenting tumour, nodule, or hyperplasia covering more than 10% of body surface, epidermis lesion, weight loss, and/or feeding/swimming troubles were sacrificed. The Kaplan–Meier method was used to calculate the probability of tumour formation and significance was established using a Mantel-Cox test (Prism 7.3; GraphPad Software).

### In situ hybridization

Adult wild-type zebrafish ovaries were fixed in 4% PFA (Electron Microscopy Science) for 24 h at 4^0^C, dehydrated, embedded in paraffin blocks and sectioned at 4 µm. To generate probes for in situ hybridisation analyses, full-length zebrafish *casc15* was amplified by PCR using Phusion DNA polymerase (ThermoScientific) from mixed-stage zebrafish cDNA. The PCR product was sub-cloned into the pGEM-T vector (Promega) and confirmed by sequencing. DIG-labelled RNA probes were generated by vector linearization and T7 in vitro transcription with the DIG-RNA labelling kit (Roche). Sections of the zebrafish ovary were deparaffined and rehydrated, fixed in 4% PFA, digested with proteinase K, acetylated and hybridized with the corresponding probe in 50% formamide, 5x SSC, 5X Denhardt’s solution, 500µg/ml salmon sperm DNA and 250 µg/ml tRNA overnight at 56^0^C. Post hybridization washes were performed in 50% formamide and 2x SSC at 56^0^C, then 2x SSC at room temperature. Sections were blocked in 10% sheep serum and incubated overnight at 37^0^C with anti-DIG-AP antibody (Roche at 1:1000). Signal detection was done using NBT/BCIP substrate (Roche).

### Histology

Entire adult zebrafish were fixed in 4% PFA (Electron Microscopy Science) for three days at 4^0^C and decalcified in 0,25mM EDTA for three more days at 4^0^C. Zebrafish were sectioned and disposed in a coronal or longitudinal orientation in histology cassettes and stored in 70% ethanol. Paraffin embedding, sectioning at 4µm and Hematoxylin-Eosin-Safran staining were performed at the Histim platform (Institut Cochin, Paris). Internal tumour Schmorl and tyrosinase staining were performed by the APEX platform (Oniris, Nantes).

### Cell line procedures

The HEK293T cell line stably expressing NanoLuc-MS2CP fusion protein^33^ was cultured in DMEM (Gibco, #41965062). 501mel cells (Lionel Larue laboratory, Institut Curie) were cultured in RPMI 1640 (Gibco, #21875091). Both medias were supplemented with 10% foetal bovine serum (Gibco, #10270106) and 1% penicillin/streptomycin (Gibco, #15140). Cells were grown at 37 °C in a humidified incubator with 5% CO2 and tested regularly for mycoplasma contamination by PCR. Plasmids or esiRNAs used for transfections were resuspended in Opti-MEM I reduced serum medium (Gibco, #31985).

### Generation of mutant and transgenic 501mel cell lines

To generate *CASC15^Δexon^*^12^ 501mel cell lines, two single guide RNAs were cloned into pSpCas9 (BB)-2A-eGFP vector (Addgene plasmid #48138). Guide RNAs were designed using CRISPick portal from the Broad Institute or using the IDT gRNA design tool^48^. 501mel cells were transfected with 500 ng of each sgRNA using Lipofectamine 2,000 Transfection Reagent (Invitrogen, #11668). 48 h after transfection, the GFP-high population of cells was collected by FACS and individual cells were seeded into each well of a 96-well plate. Sorted cells were genotyped by PCR and DNA sequencing.

To generate the HA-tagged DDX50 cell lines, an HA-PreScission-6xHis tag was introduced at the N-terminus of DDX50 and DUSP12. The tag flanked by ∼400 nt homology arms was cloned into the pUC57 vector. 501mel cells were transfected with DDX50 or DUSP12 HA-targeting plasmids together with the pSpCas9(BB)-2A-Puro vector (Addgene plasmid #62988) containing a gRNA sequence targeting each of the proteins. Lipofectamine 2000 Transfection Reagent (Invitrogen, #11668) was used for transfections. Transfected cells were selected with 1 µg/mL of puromycin for 48 h. For each transfection, individual clones were expanded and genotyped for the integration of the tag at the correct genomic location by PCR and DNA sequencing.

Genomic DNA was extracted by lysing the cells with lysis buffer (10 mM Tris-HCl pH 7.5, 10 mM EDTA, 10 mM NaCl, 0.5% SDS) supplemented with 0.1mg/ml proteinase K (Roche, #03115828001) at 55 °C for 3 h, followed by ethanol precipitation and resuspension in nuclease-free water. All relevant oligonucleotides and tag sequences are listed in Supplementary Table 2.

### Cellular assays

For the proliferation assay, ∼30,000 cells were plated in a 35 mm dish at day 0 and counted (Vi-cell XR, Beckman Coulter) each day until day 5. For the clonogenicity assay, 35 mm dishes were seeded with ∼500 cells. After 10 days of growth, colonies were fixed with 4% PFA, stained with Crystal violet in 10% ethanol and counted.

For the invasion assay, transwell chambers with 8 µm pore size (Falcon, #353097) were coated with 400 µg/ml of GFR Matrigel matrix (Corning, #354230). A cell suspension of ∼200.000 cells in serum-free media was added to the Matrigel-coated upper chamber and FBS-supplemented media was added to the lower chamber as chemoattractant. Cells were allowed to invade for 24 h, then fixed with 4% PFA, and stained with Crystal violet in 80% methanol, 10% formaldehyde, and counted.

For the wound-healing assay, 30,000 cells were seeded on each side of silicone inserts (Ibidi, #80209). After 24 h, inserts were removed, and migration was recorded by imaging cells every hour for a period of 24 h using an IncuCyte Live Cell Imaging System (Essen, Bioscience) with a 4x magnification. Wound closure was evaluated by measuring the size of the wound with ImageJ software using an edge-detection macro. All experiments were performed in three biological replicates (independently generated cell lines) for each genotype in technical triplicates.

### esiRNAs

Protein depletion was achieved using esiRNAs (Eupheria Biotech) targeting mRNAs coding for indicated proteins. esiRNAs targeting *RLuc* were used as a negative control. For the wound-healing assay, cells in each of the two sides of the IBIDI inserts were transfected with 0.05 µg of esiRNAs and 0.5 µl of Lipofectamine 2,000 Transfection Reagent (Invitrogen, #11668) 24 h before the start of the migration assay. To assess esiRNA knockdown efficiencies, both total RNA and whole cell lysates were extracted 24 h post-transfection followed by qRT-PCR and Western blot analysis, respectively.

### incPRINT expression constructs

To generate RNA-MS2 expression constructs, PCR-amplified full-length, truncated and antisense fragments of zebrafish and human *CASC15* were cloned into the BstBI restriction site of the pCDNA3.1 plasmid containing MS2 stem loops^33^ using Gibson assembly (New England Biolab, #E2621). Human *CASC15* was PCR-amplified from Human Brain Total RNA (ThermoFisher #AM7962); zebrafish *casc15* was PCR-amplified from zebrafish embryo cDNA. All relevant oligonucleotides used for *CASC15* fragment amplification and cloning are listed in Supplementary Table 2.

### incPRINT

incPRINT was carried out as previously described^33^. Briefly, the HEK 293T stable cell line expressing NanoLuc luciferase fused to MS2CP was co-transfected with plasmids encoding for RNA-MS2 and 3xFLAG-tagged protein using 300ng and 600ng of plasmids, respectively. Two days post-transfection, cells were PBS-washed twice, lysed and DNase treated. Lysates were transferred into an anti-FLAG coated 384-well plate and incubated for 3 h. After seven washes, luciferase substrate was added to the plates and luminescence was read, and subsequent ELISA was performed using an HRP-conjugated anti-FLAG antibody and its substrate. Interaction values between each tested RNA-MS2 and a test protein were defined as the average between the two luminescence replicates. To compare the interaction intensities between high-throughput experiments using different RNA-MS2 transcripts, raw luminescence intensities were normalized as previously described^33^. Briefly, median interaction scores of a set of proteins that interact with MS2 stem loops alone were used as a ratio for all raw luminescence intensities measured for the corresponding RNA-MS2^33^. We considered protein interactors as shared between zebrafish and human *CASC15* transcripts when the normalized score was above 1.5 for both transcripts and among the top interactors for at least one RNA (above 2.2 for human *CASC15* and 2.0 for zebrafish *casc15)* (**Supplementary Table 1**).

### RNA immunoprecipitation (RIP)

∼3,5 million 501mel cells were seeded in a 15 cm dish, transfected the following day with 15 µg of plasmid to exogenously express HA-tagged proteins and incubated for 48 h. For the DDX50^+/HA^ 501mel cell line, nine million cells were seeded and incubated for 24 h. Cells were washed once with ice-cold 1x PBS and UV-crosslinked using 800 mJ/cm^2^, at 254 nm (Stratalinker, Stratagene). Cells were collected and lysed for 10 min on ice in 200 µl of RIPA buffer (50 mM Tris-HCl pH 8, 150 mM NaCl, 1% Triton X-100, 0.5% sodium deoxycholate, 0.1% SDS) with freshly added 1x cOmplete EDTA-free protease inhibitor cocktail (Roche, #11873580001) and 100U/ml of RNAseOUT. Cell lysates were sonicated using a Bioruptor (Diagenode) (6 cycles; 15s on, 30s off, M) and centrifuged for 10 min at 15,000x g. Supernatant were collected and diluted in RQ1-HENG buffer to 1 ml, and 1% was saved as total cell lysate input for western blot analysis. 50 µl of anti-HA bead slurry (Pierce, #88836) were washed twice with 1ml of RQ1-HENG buffer and incubated with the cell lysate for 2h at 4 °C. Following the incubation, beads were washed four times with 1 ml of RQ1-HENG. Proteins were then eluted from the beads in elution buffer (10 mM Tris pH 8, 1 mM EDTA, 1% SDS) for 2 min at 90^0^C, followed by proteinase K (Roche, #03115828001) treatment for 30 min at 37^0^C. 20% of each eluate was collected for further analysis by Western blot analyses. RT-qPCR data was normalized to *ß-ACTIN* and RNA enrichment in the eluate was displayed relative to input.

### Patients and Samples

Samples of 94 primary melanomas and 14 metastatic melanomas, from patients managed at Institut Curie Hospital (France) as well as cell lines have been analysed. All patients who entered our institution were informed that their tumour samples might be used for scientific purposes and had the opportunity to decline. Since 2007, patients entering our institution have given their approval also by signed informed consent. This study was approved by the local ethics committee of Institut Curie Hospital. The samples were immediately stored in liquid nitrogen until RNA extraction. A tumour sample was considered suitable for our study if the proportion of tumour cells exceeded 70%.

Fifty specimens of normal tissues of the skin were used as sources of normal RNA.

### qRT-PCR analysis

Total RNA from zebrafish samples, 501mel cells or RIP experiments was isolated using TRIzol (Invitrogen, #15596) according to manufacturer’s instructions. Extraction was followed by DNAse treatment (TURBO DNA-free, Invitrogen #AM2238) and ethanol precipitation. 200 ng of total RNA isolated from RIP experiments and 900 ng of total RNA isolated from all other samples were reverse-transcribed using SuperScript IV kit with oligo-dTs (Invitrogen, #18091), followed by qPCR using SYBR Green (Applied biosystems, #4767659) according to manufacturer’s instructions. *ß-ACTIN*, *TBP* or *eif1α* were used as reference genes. The number of biological replicates is indicated in figure legends; each biological replicate was run with four technical replicates. All relevant oligonucleotides are listed in **Supplementary Table 2**.

Total RNA extraction from human samples, cDNA Synthesis and PCR reaction conditions have been performed as previously described^49^. Human samples were analysed for expression of selected genes using real-time reverse transcription-PCR (RT-PCR); Quantitative real-time RT-PCR was performed as previously described^49^. Quantified levels of TATA box-binding protein (TBP) transcripts were used as the endogenous RNA control, and each sample was normalized on the basis of its TBP content. Results, expressed as N-fold differences in *CASC15* gene expression relative to the *TBP* gene, and termed “N*CASC15*”, were determined as N*CASC15* = 2^DCtsample^, where the DCt value of the sample was determined by subtracting the average Ct value of the *CASC15* gene from the average Ct value of the *TBP* gene. The N*CASC15* values of the samples were subsequently normalized such that the mean ratio of the normal skin samples would equal a value of 1. All relevant oligonucleotides are listed in **Supplementary Table 2**.

### Western blot analysis

Protein samples collected from RIP were denaturated at 95^0^C for 5 min, resolved on 4–12% NuPAGE gels and transferred to a nitrocellulose membrane (Amersham Protran, GE Healthcare Life Sciences, #10600002) using a wet-transfer system. After blocking with 5% non-fat milk in TPBS (PBS with 0.1% Tween-20) for 1 h at room temperature, the membranes were incubated overnight at 4^0^C with the anti-HA antibody (Cell Signaling, #C29F4, 1:1,000 dilution). Membranes were washed twice with TPBS and incubated for 1h at room temperature with the secondary antibody conjugated with horseradish peroxidase (Promega, #W401B, 1:5,000 dilution) followed by three more washes. Signal was detected using (Amersham Protran, GE Healthcare Life Sciences, #RPN2232) on the Chemidoc MP imaging system (BioRad). Full blots are provided as a Source Data file.

### RNA-Seq data processing and differential expression analyses

Following RNA extraction, 1 µg of total RNA was used for RNA-Seq library preparation using Illumina Stranded mRNA prep Ligation according to the manufacturer’s recommended protocol. 100bp paired-end reads were generated using the Illumina NovaSeq platform. Adapter sequences were trimmed from raw reads using TrimGalore! (v0.6.7). Trimmed reads were first mapped to the complete human rRNA sequence with bowtie (v1.3.0). Unmapped reads were subsequently mapped with STAR (v2.7.6a) to the human reference genome (hg19/GRCh37) and gene counts were generated using --quant_mode GeneCounts. Strandedness (reverse) was confirmed using RSeQC (v4.0.0). Gene name annotations were obtained from gencode (v19).

RNA-seq data were processed with the Institut Curie RNA-seq pipeline (v3.1.8) (La Rosa Philippe, Fabrice Allain Philippe Hupé, Nicolas Servant. (2022). bioinfo-pf-curie/RNA-seq: v3.1.8 (v3.1.8). Zenodo. https://doi.org/10.5281/zenodo.7446922). Genes were filtered to include those with CPM > 0.5 in 2 or more samples. Raw count data was transformed to log2-CPM and normalized with the TMM method using edgeR (v3.54.2; Robinson 2010 and McCarthy 2012). A linear model was fit to the normalized expression values for each gene and empirical Bayes statistics were computed for each comparison with limma (v3.40.2; Ritchie, 2015). Differentially expressed genes for each KO versus WT were identified from the linear fit after adjusting for multiple testing and filtered to include those with FDR < 0.05 and absolute logFC > 1.

Enrichments analyses were performed with ClusterProfiler (v4.0.5)^50^ using an adjusted p-value (FDR) of 0.05 with all genes used in the differential expression analysis as background.

### RNA motif analysis

The *k*-mer content analyses of human *CASC15* was performed using SEEKR algorithm (https://app.med.unc.edu/seekr/home). All human lncRNAs (Gencode, V40) were used as normalization set and transformation was performed pre-standardization. For vertebrate-conserved RNA-motif discovery, lncLOOM algorithm was used as described^21^ using *CASC15* species from different vertebrates. The lncRNA sequences used for the analysis are listed in **Supplementary Table 3**.

### Data Availability

RNA-Seq data generated in this study are available in the NCBI Gene Expression Omnibus (http://www.ncbi.nlm.nih.gov/geo/). The accession number for the sequencing data reported in this paper is under accession NCBI GEO: GSE235839. No previously unpublished algorithms were used to generate the results.

## AUTHOR CONTRIBUTIONS

X.S.C. and P.L. contributed to the design, execution, and analyses of most experiments. X.S.C. prepared figures and wrote the first version of the manuscript. C.J.R. and I.U. performed computational analyses of *CASC15* sequences including LncLOOM. L.C. contributed to the execution of cellular assays, *CASC15* sequence conservation analyses and figure preparation. D.Z. and N.S. performed computational analyses including RNA-seq data analyses. S.V. and I.B. performed expression analysis on human melanoma cell lines and patient biopsies. M.L. and T.L. performed histo-pathological analysis of the internal zebrafish tumours. M.P. and L.D. assisted with *in vivo* and cellular experiments. A.S. conceived and supervised the study, acquired funding and wrote the manuscript.

## ACKNOWLEDGEMENTS

We thank Adam Hurlstone for providing the pDEST-Tol2-*mitfa:NRAS^G12D^* plasmid and for his advice on zebrafish melanoma assays. We thank Lionel Larue for providing cell lines used in the study and for his advice on cellular assays. We also thank all members of the Shkumatava laboratory for stimulating discussions. This work was supported by funding from the Tandem Weizmann–PIC3i Curie 2019-2021 grant, Fondation pour la Recherche Médicale (EQU202003010550), LABEX DEEP (ANR-11-LABX-0044_DEEP, ANR-10-IDEX-0001-02_PSL), Franco-Israeli Projects of Research Collaboration CNRS-MoST and Institut Curie. X.S-C. was supported by doctoral fellowships from the IC-3i international PhD program (H2020 program under the Marie Skłodowska-Curie grant agreement no. 666003) and Fondation pour la Recherche Médicale. P.L. was supported by doctoral fellowships from the PSL University and La Ligue nationale contre le cancer. High-throughput sequencing has been performed by the ICGex NGS platform of the Institut Curie supported by the grants ANR-10-EQPX-03 (Equipex) and ANR-10-INBS-09-08 (France Génomique Consortium) from the Agence Nationale de la Recherche (“Investissements d’Avenir” program), by the Canceropole Ile-de-France and by the SiRIC-Curie program - SiRIC Grant (INCa-DGOS-4654). The imaging experiments were performed in the Cell and Tissue Imaging Facility/ UMR3215 (PICT-IBiSA) of Institut Curie, member of the French National Research Infrastructure France-BioImaging (ANR-10-INBS-04).

## SUPPLEMENTARY FIGURES

**Supplementary Figure 1:**
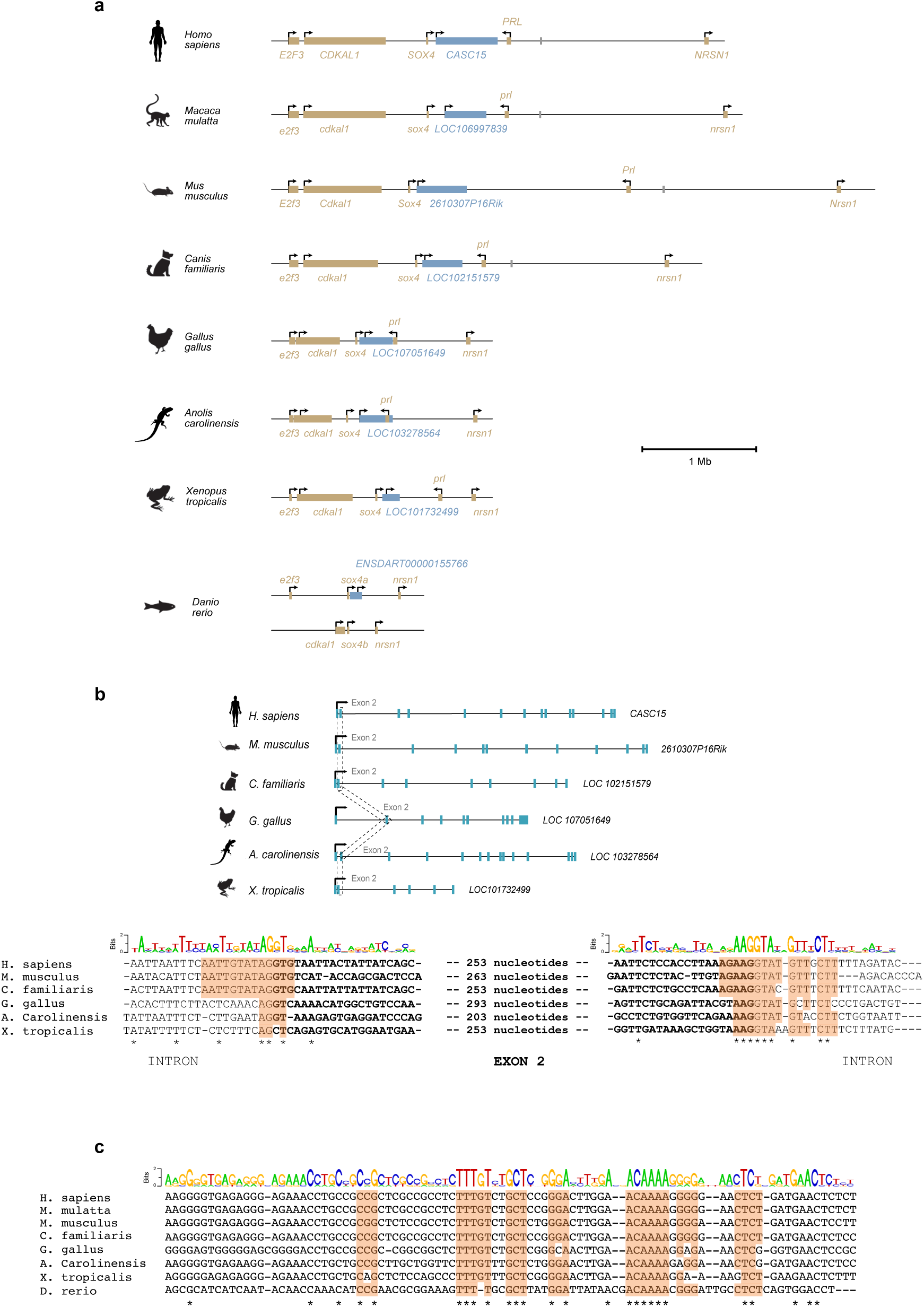
Synteny and sequence conservation of the *CASC15* locus, related to Figure 1. **(a)** Conservation of the order and transcriptional orientation of genes in the *SOX4/CASC15* locus in different species. Transcriptional homologs of the lncRNA *CASC15* are indicated in blue. **(b)** Top panel, genomic locus of *CASC15* in different vertebrates. Representative isoforms of *CASC15* are shown. The dotted line indicates the location of conserved splice junctions. Bottom panel, conservation plot relative to the human *CASC15* locus. The sequence logo based on four mammalian homologous sequences is shown above. Motifs conserved between human and tetrapods exon 2 splice junctions are highlighted in orange. **(c)** Conservation plot relative to the human *CASC15* locus. The sequence logo based on four mammalian homologous sequences is shown above. Motifs conserved between human and zebrafish *CASC15* transcriptional homologs are highlighted in orange.

**Supplementary Figure 2:**
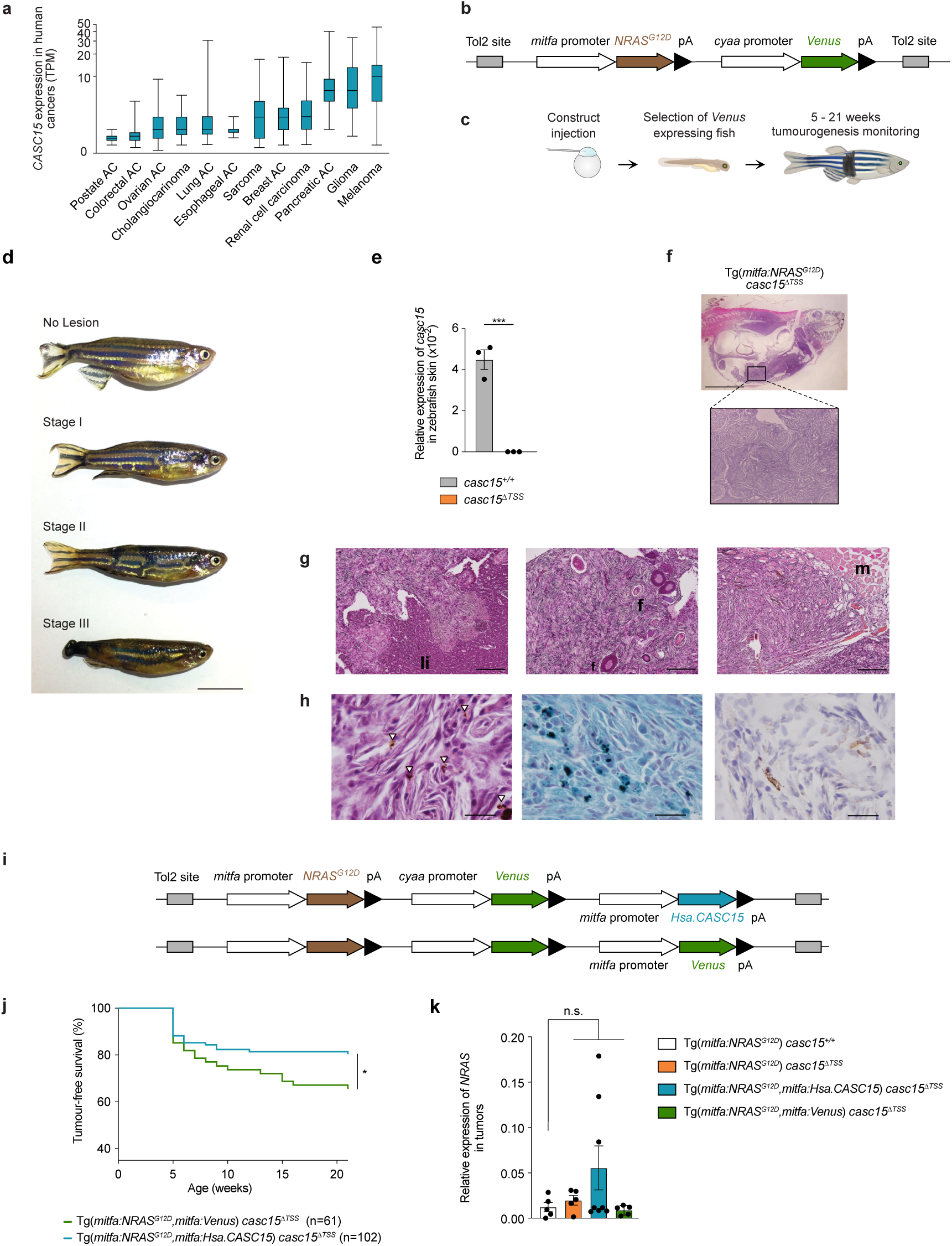
Experimental strategy for testing the role of *casc15* in melanoma formation, related to Figure 2. **(a)** Expression of human *CASC15* across different human cancers from the Pan Cancer Analysis of Whole Genome (PCAWG) project. TPM, transcripts per million. AC, Adenocarcinoma. **(b)** Schematic representation of the NRAS^G12D^ expression construct used to induce melanoma in zebrafish. pA, polyadenylation signal. **(c)** Experimental set-up to analyse melanoma tumorigenesis in zebrafish. **(d)** Representative images of different stages of melanoma formation in wild-type zebrafish expressing NRAS^G12D^ driven by the *mitfa* promoter at seven weeks post-fertilization (wpf). Arrows indicate pigmentation deficits. Scale bar represents 1cm. **(e)** qRT-PCR analysis of *casc15* expression in wild-type and *casc15^ΔTSS^* zebrafish skin. *eef1α1* was used as a reference gene. Data presented as mean ± s.e.m.; unpaired t-tests: \*\*\**P* < 0.001 **(f)** Haematoxylin and eosin staining of whole-body sagittal sections of *casc15^ΔTSS^*zebrafish expressing NRAS^G12D^ driven by the *mitfa* promoter at 19 weeks post-fertilization (wpf). Scale bar represents 1cm. **(g)** Haematoxylin and eosin staining of infiltrating internal tumour sections of *casc15^ΔTSS^* zebrafish expressing NRAS^G12D^ driven by the *mitfa* promoter at 19 weeks post-fertilization (wpf), presenting a stage III tumour. Scale bar represents 200 µm. Li, liver parenchyma; f, follicle; m, skeletal muscles layers. **(h)** Haematoxylin and eosin, Schmorl and anti-tyrosinase staining of internal tumour sections of *casc15^ΔTSS^* expressing NRAS^G12D^. Arrow heads indicate intracytoplasmic brown pigments typical of melanocytic granules. Scale bar represents 20 µm. **(i)** Schematic representation of constructs used for rescue experiments. pA, polyadenylation signal. **(j)** Kaplan-Meier plot of melanoma-free period for wild-type and *casc15^ΔTSS^*zebrafish expressing either the *NRAS^G12D^ + Venus* or *NRAS^G12D^ + human CASC15* transgenes, driven by the *mitfa* promoter. Data are analysed using Mantel-Cox test: \**P* < 0.05. **(k)** qRT-PCR analysis of *NRAS^G12D^* mRNA expression in tumours isolated from zebrafish expressing either *NRAS^G12D^* alone, *NRAS^G12D^* + human *CASC15* or *NRAS^G12D^* + *Venus* transgenes, driven by the *mitfa* promoter; wild-type zebrafish are indicated as *casc15^+/+^*. *eef1α1* was used as a reference gene. Data presented as mean ± s.e.m.; unpaired t-tests: not significant (ns).

**Supplementary Figure 3:**
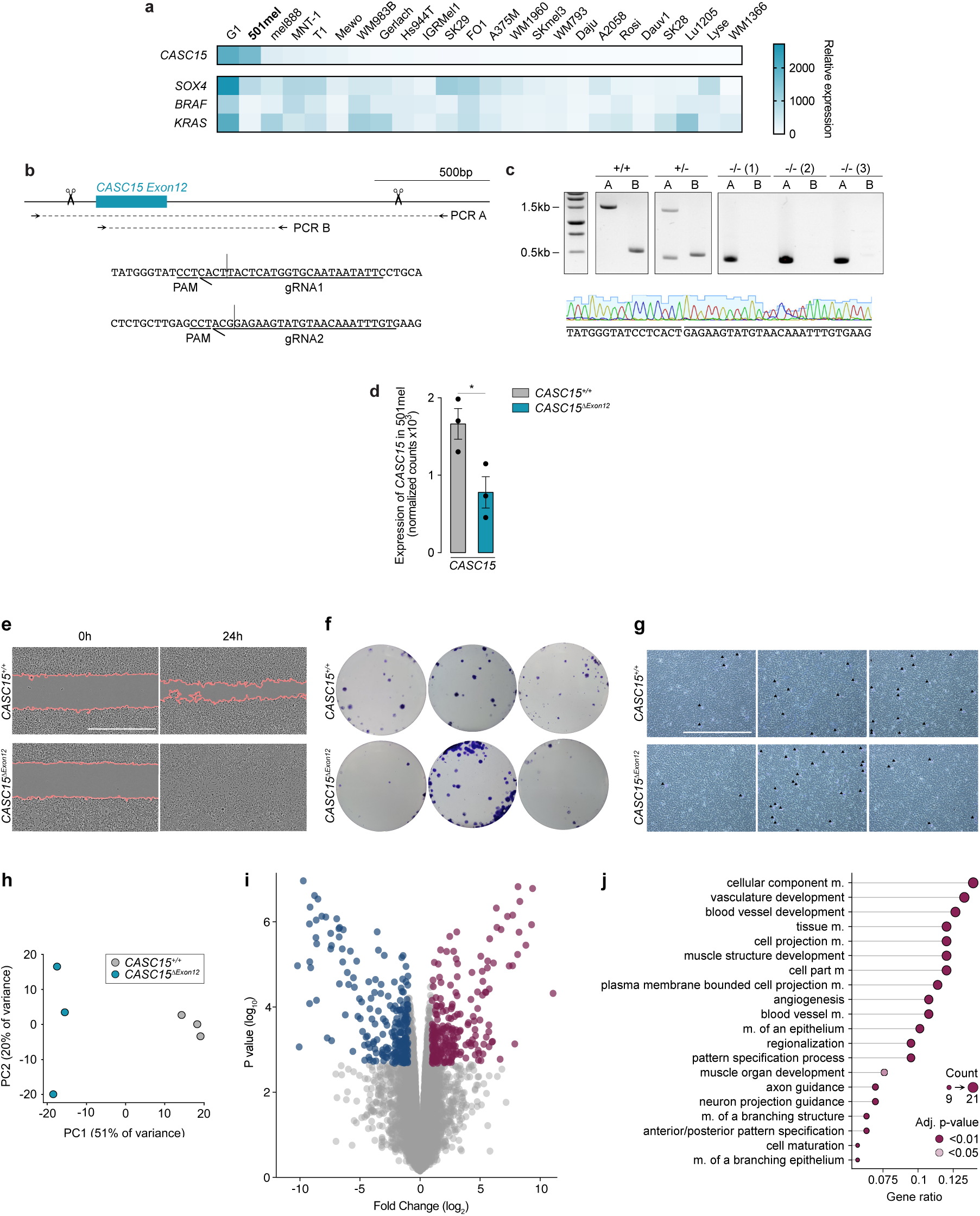
Experimental strategy for generation of the *CASC15* null allele in 501mel cells, related to Figure 3. **(a)** Heatmap of human *CASC15* expression levels analysed by qRT-PCR in indicated melanoma cell lines. *TBP* was used as a reference gene. **(b)** Targeting and genotyping strategy for the deletion of exon 12 of the *CASC15* locus in 501mel cells. Scissors indicate the location of gRNAs; arrows indicate the location of primers used for genotyping of independently derived cell lines. gRNA sequences are indicated below and primer sequences are listed in Supplementary Table 2. **(c)** Genotyping results for the *CASC15^Δexon^*^12^ 501mel cells. Top panel, representative agarose gel images showing amplicons from the unedited *CASC15* locus (+/+), heterozygous (+/-) and homozygous (-/-) deletion alleles of exon 12 of *CASC15*. Bottom panel, representative sequencing trace of an amplicon from a homozygous *CASC15^Δexon^*^12^ mel501 cell line. **(d)** *CASC15* expression levels detected by RNAseq: * = *P* < 0.05 **(e)** Representative images of the wound-healing assay. Scale bar represents 1 mm **(f)** Representative images of the colony formation assay. Scale bar represents 1 mm **(g)** Representative images of the invasion assay. Scale bar represents 500 µm **(h)** PCA plot of regularized log-transformed RNA-Seq counts from *CASC15^+/+^* and *CASC15^Δexon^*^12^ 501mel cell lines. Each dot represents a biological replicate. **(i)** Volcano plot showing changes in gene expression in *CASC15^Δexon^*^12^ 501mel cells compared to *CASC15^+/+^* cells. Coloured points indicate genes with enrichment >2 fold and FDR adjusted *P* < 0.05. **(j)** Lollipop graph with biological process enrichment of upregulated genes.

**Supplementary Figure 4:**
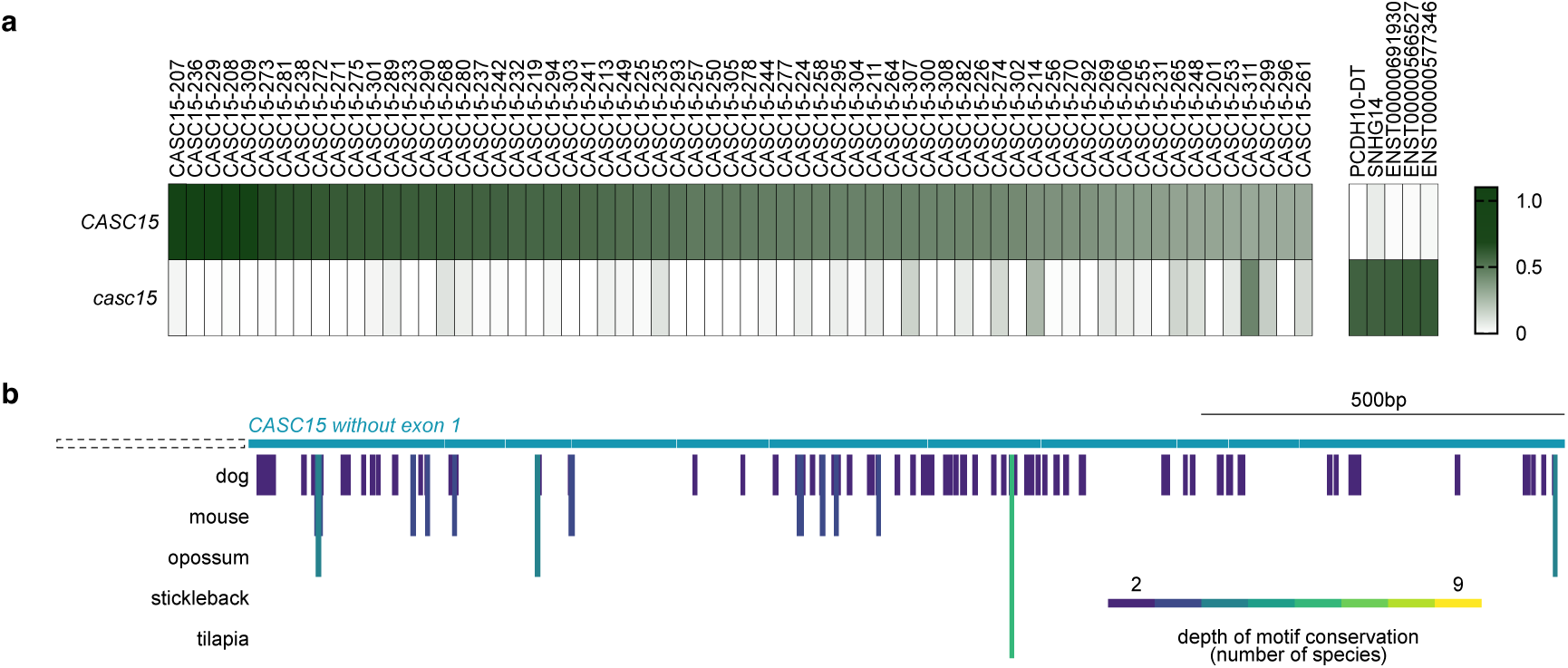
Analyses of conserved RNA motifs using SEEKR and LncLOOM algorithms with less stringent parameters, related to Figure 4. (**a**) *k*-mer content analysis between indicated *CASC15* orthologs and the human transcriptome using SEEKR algorithm. Left panel shows the top 65 transcripts most similar to human *CASC15;* right panel shows the top five transcripts most similar to zebrafish *casc15*. *k*=5; all human lncRNAs were used as normalization set. (**b**) Distribution of short conserved RNA motifs within exons of *CASC15* identified by the LncLOOM algorithm. Exon 1 was removed from all analysed *CASC15* transcripts. Identified elements are represented by vertical bars color-coded by conservation depth. Vertebrate species that have identified motives are indicated on the left. The human *CASC15* transcript was used as anchor.

**Supplementary Figure 5:**
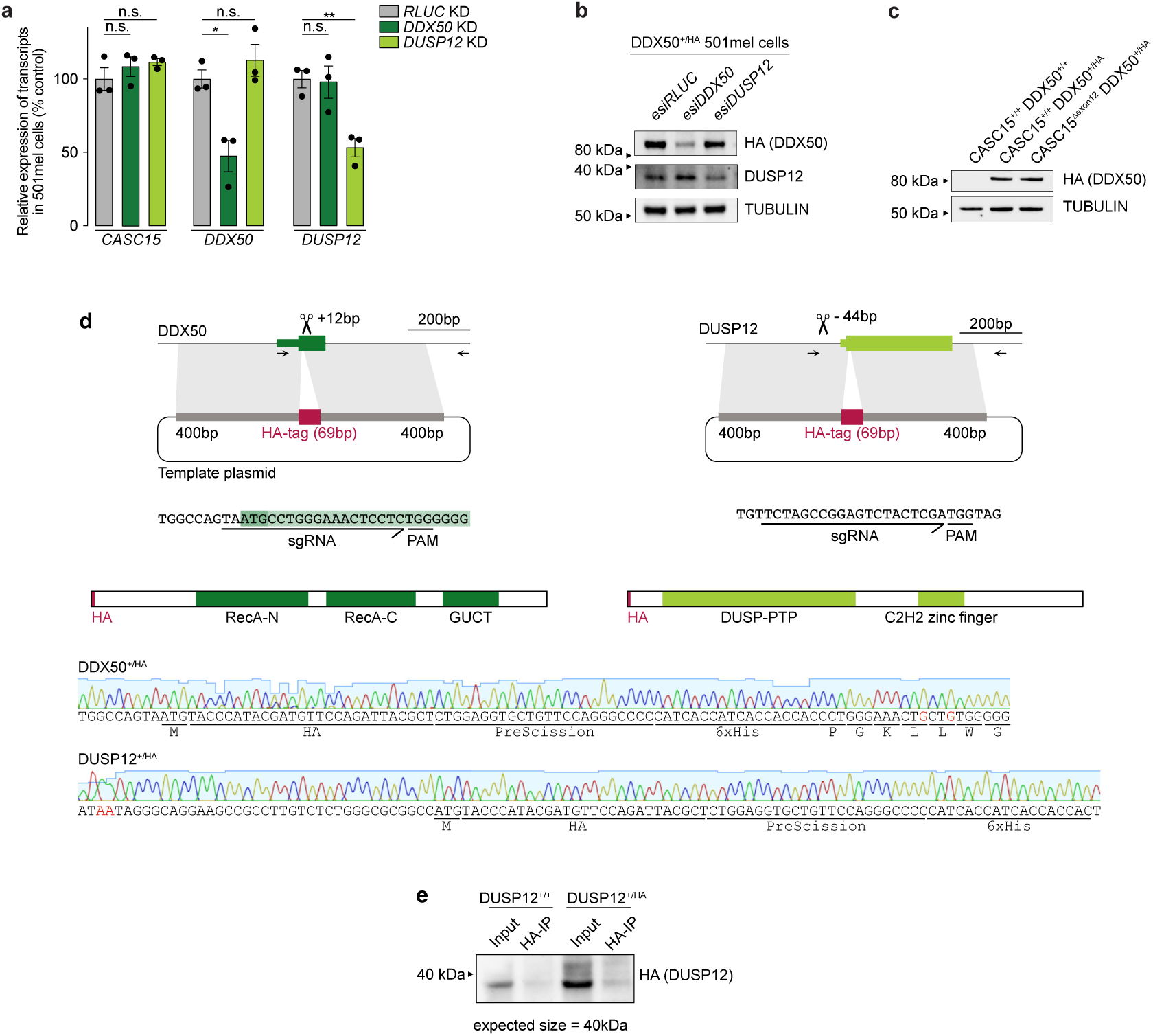
Quantification of esiRNA efficiency and experimental strategy for the endogenous protein tagging in 501mil cells, related to Figure 5. **(a)** qRT-PCR analyses of *CASC15*, *DDX50* and *DUSP12* levels upon their respective esiRNA-mediated depletion in 501mel cells. *ß-ACTIN* was used as reference gene and results were normalized to the corresponding *RLUC* negative control. Data from three transfection replicates are presented as mean ± s.e.m; unpaired t-test; * *P* < 0.05; ** *P* < 0.01; not significant (ns). **(b)** Western blot analysis of the efficiency of the esiRNA-mediated protein knock-down using anti-HA and anti-DUSP12 antibodies. *RLUC* esiRNA was used as a control. TUBULIN was used as a loading reference. **(c)** Western blot analysis of the endogenous DDX50 protein levels of the engineered cell lines on indicated *CASC15^+/+^* or *CASC15^Δexon^*^12^ backgrounds, using anti-HA antibody. **(d)** Schematic representation of the DDX50 locus and design of the sgRNA targeting the for the insertion of the HA-tag into the 5′ terminus of the DDX50 locus. Top panel, schematic of the template vector and CRISPR-mediated homologous recombination targeting strategy. Genotyping primers are indicated with arrows. Middle panel, sgRNA and the protein sequence scheme tagged with the HA epitope tag. Bottom panel, representative sequencing traces of the tagged region of heterozygous cell lines. In red, nucleotides changed to inactivate the template without changing the amino acid sequence. GUCT = ‘Gu’ -patient C-terminal domain. All relevant oligonucleotides and tag sequences are listed in Supplementary Table 2. **(e)** Representative western blot on input or HA-immunoprecipitated samples isolated from *DUSP12^+/+^* or *DUSP12^+/HA^* 501mel cells, using anti-HA antibody.

